# Structural basis of reiterative transcription from the *pyrG* and *pyrBI* promoters by bacterial RNA polymerase

**DOI:** 10.1101/732214

**Authors:** Yeonoh Shin, Mark Hedglin, Katsuhiko S. Murakami

## Abstract

Reiterative transcription is a non-canonical form of RNA synthesis by RNA polymerase in which a ribonucleotide specified by a single base in the DNA template is repetitively added to the nascent RNA transcript. We previously determined the X-ray crystal structure of the bacterial RNA polymerase engaged in reiterative transcription from the *pyrG* promoter, which contains 8 poly-G RNA bases synthesized using 3 C bases in the DNA as a template and extends RNA without displacement of the promoter recognition σ factor from the core enzyme. In this study, we determined a series of transcript initiation complex structures from the *pyrG* promoter using soak trigger freeze X-ray crystallography. We also performed biochemical assays to monitor template DNA translocation during RNA synthesis from the *pyrG* promoter and *in vitro* transcription assays to determine the length of poly-G RNA from the *pyrG* promoter variants. Structures and biochemical assays revealed how the RNA transcript from the *pyrG* promoter is guided toward the Rifampin-binding pocket then the main channel of RNA polymerase and provided insight into RNA slippage during reiterative transcription of the *pyrG* promoter. Lastly, we determined a structure of a reiterative transcription complex at the *pyrBI* promoter and revealed an alternative mechanism of RNA slippage and extension requiring the *σ* dissociation from the core enzyme.

**SIGNIFICANCE STATEMENT:** RNA polymerase synthesizes multiple bases of RNA using a single base of the template DNA due to slippage between RNA transcript and template DNA. This noncanonical RNA synthesis is called “reiterative transcription,” playing several regulatory roles cellular organisms and viruses. In this study, we determined a series of X-ray crystal structures of a bacterial RNA polymerase engaged in reiterative transcription and characterized a role of template DNA during reiterative transcription by biochemical assays. Our study revealed how RNA slips on template DNA and how RNA polymerase and template DNA determine length of reiterative RNA product. We also provide insights into the regulation of gene expression using two alternative ways of reiterative transcription.

## INTRODUCTION

Non-canonical form of transcription called “reiterative transcription” (also known as transcript slippage) regulates gene expression (1, 2). Unlike canonical transcription in which RNAP simply copies the DNA sequence to RNA, RNAP adds extra bases to the RNA during reiterative transcription. This is due to repetitive addition of the same nucleotide to the 3’ end of a nascent RNA while RNA slips upstream on the template DNA (1–6). Since first proposed in the early 1960s in the *Escherichia coli* RNAP transcription (3), reiterative transcription has been discovered and characterized not only in cellular RNAPs from bacteria to human but also in virus RNAPs (7–15).

The *pyrG* gene in *Bacillus subtilis* encodes CTP synthetase and its expression is regulated by CTP-dependent reiterative transcription (**Fig. S1**) (6). The initially transcribed region (ITR) of *pyrG* (5’-<u>G</u>GGCTC on the non-template DNA and 3’-<u>C</u>CCGAG on the template DNA, the transcription start site is underlined) contains a slippage-prone homopolymeric DNA sequence followed by a base that determines the fate of RNA extension, either canonical or reiterative, depending on the amount of CTP. In the presence of a high concentration of CTP, RNAP transcribes RNA without slippage (5’-GGGCUC) and continues until an attenuator sequence, which forms the transcription termination hairpin thereby eliminating *pyrG* expression (**Fig. S1, left**). On the other hand, when CTP is limited, right after 5’-GGG-3’ RNA is synthesized, RNAP starts reiterative transcription and inserts up to 10 extra G bases to the nascent RNA before returning to canonical transcription (5’-GGGGnCUC, n=1∼10), resulting in the formation of an anti-termination hairpin with the pyrimidine-rich sequence in the 5’ part of the attenuator, thereby allows expression of the *pyrG* gene (**Fig. S1, right**).

Another well-known example of conditional reiterative transcription is UTP-sensitive regulation of transcript initiation at the *pyrBI* operon of *E. coli* (16). The *pyrBI* ITR (5’-<u>A</u>A*TTT*G, non-template DNA, transcription start site is underlined) contains a slippage prone sequence (italicized) (**Fig. S2**), where transcript slippage produces transcripts with the sequence 5’- AAUUUn (where n=1 to >100) (16). In contrast to the regulation of the *pyrG* promoter where reiterative transcription eventually switches to canonical transcription to express the *pyrG* operon (**Fig. S1**), there is no switch to canonical transcription from reiterative transcription at the *pyrBI* promoter. Instead, the reiterative transcripts are released from the transcript initiation complex (**Fig. S2, left**) (2).

Previously, we reported the X-ray crystal structure of the reiterative transcription complex (RTC) from the *pyrG* promoter, which was prepared by *in crystallo* RNA synthesis in the presence of GTP in a 30 min reaction within the bacterial RNAP and *pyrG* promoter DNA complex crystal (RTC-30’) (17). The structure represented the final stage of reiterative transcription, revealed the presence of 8-mer poly-G RNA and showed that 3 bases at the 3’ end form a DNA/RNA hybrid and a fourth base from the 3’ end of RNA (−4G) fits into the Rifampin (RIF) binding pocket of the *β* subunit of RNAP. These features allow RNA to detour from the dedicated RNA exit channel and extend toward the main channel of the enzyme without displacement of the σ factor. The 3’ end of RNA is in a post-translocated state (i.e., in the *i* site), forming a base pair with template DNA residue +3C, whereas the +4G base is positioned at the *i+*1 site, poised for incoming CTP to extend the nascent RNA by canonical transcription. The structure revealed an unexpected RNA extension pathway during reiterative transcription; however, several questions remain to be answered such as how RNA slips on template DNA and how the 5’ end of RNA is guided toward the main channel of RNAP.

In this study, we further study the mechanism of reiterative transcription from the *pyrG* promoter by structural and biochemical approaches. We determined a series of X-ray crystal structures of transcript initiation complexes containing 2-, 3- and 4-mer RNAs. Additionally, we determined a series of structures with *pyrG* promoter variants containing base substitution at the template DNA −1 position and revealed a role of the template DNA base for guiding RNA toward the RIF-binding pocket. Lastly, we investigate the reiterative transcription from *pyrBI* promoter by structural and biochemical studies to reveal an alternative way of RNA extension compare with the *pyrG* promoter transcription.

## RESULTS

### Capturing transcript initiation complexes by time-dependent soak-trigger-freeze X-ray crystallography

We applied time-dependent soak-trigger-freeze X-ray crystallography (18) to determine a series of structures representing the transcript initiation complex from the *pyrG* promoter. We previously demonstrated that *Thermus thermophilus* σ^A^ RNAP holoenzyme is proficient at reiterative transcription from the *B. subtilis pyrG* promoter both *in vitro* and *in crystallo* (17). The crystals of *T. thermophilus* RNAP and DNA complex containing the *pyrG* promoter sequence (**Table. S1**) were soaked into a cryo-solution containing GTP to trigger RNA synthesis *in crystallo*. The reaction was stopped by freezing crystals at different time points (from 1 min to 2 hours, **Fig. 1A**) and the structures were determined by molecular replacement (**Table. S4**). Each structure present here shows electron density corresponding to in *crystallo* synthesized RNA, allowing us to monitor extension of poly-G RNA. The length of RNA increases as the crystal soaks in the GTP solution (**Fig. 1B**).

**Fig. 1.**
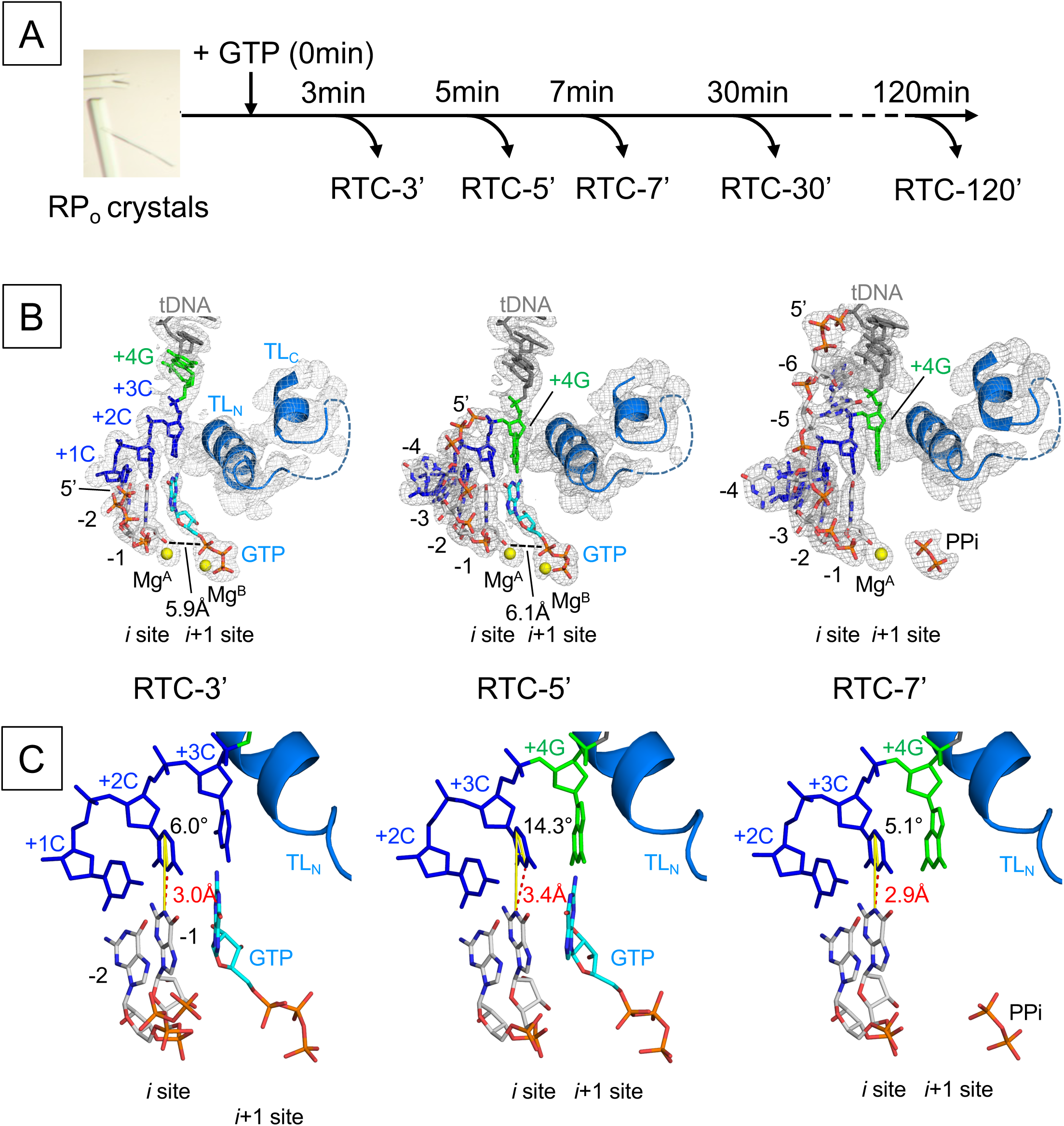
The structures of reiterative transcription complex from the *pyrG* promoter. **A)** Experimental scheme of the RTC preparation by time-dependent soak-trigger-freeze X-ray crystallography. **B)** Structures of the RTC-3’ (left), RTC-5’ (middle) and RTC-7’ (right). RNA, tDNA and incoming GTP are shown as stick models, the Mg ions bound at the active site of RNAP are shown as yellow spheres, and the RNAP trigger loop is depicted as a ribbon model. The 2Fo–Fc electron densities for RNA, template DNA and trigger loop are shown (gray mesh, 1.5 σ). **C)** Analysis of the DNA and RNA base parings at the *i* site of RNAP active site. The distance between N1 atom (RNA G base) and N3 atom (tDNA C base) is shown as a red dashed line. The angle of the DNA and RNA base pairing (from N1(RNA G base) to C5(tDNA C base) to N3(tDNA C base)) is shown as yellow lines.

After 3 min of GTP soaking (RTC-3’, **Fig. 1B, left**), RNAP synthesizes 2-mer RNA and the 3’ end of the RNA is in a post-translocated state, forming a base pair with +2C template DNA (tDNA). Hereafter, RNA residues are counted −1, −2, −3 from the 3’ end. The +3C tDNA is positioned at the *i*+1 site and forms a base pair with an incoming GTP. The α-phosphate of GTP is 5.9 Å away from the 3’-OH of RNA and the trigger loop is in the open conformation, indicating that the GTP is positioned at the pre-insertion site. The structure of RTC after 4 min of GTP soaking (RTC-4’, data not shown) was the same as the structure of the RTC-3’.

After 5 min of GTP soaking (RTC-5’, **Fig. 1B, middle**), RNA extends to 4-mer and the 3’ end of the RNA is in a post-translocated state, forming a base pair with +3C tDNA, while +4G tDNA is positioned at the *i*+1 site. The first three bases of RNA are synthesized by canonical transcription whereas the 4^th^ base is added by reiterative transcription. The 5’ end of 4-mer RNA (−4G) is inserted into the RIF-binding pocket, as observed in the previously published RTC structure (RTC-30’) containing 8-mer RNA (17), indicating that the 5’-end of RNA base fits in the RIF-binding pocket followed by extension toward the main channel of RNAP. In the RTC-5’, incoming GTP bound at the *i*+1 site forms a mismatch with the +4G tDNA and the trigger loop is in the open conformation.

At 7 min (RTC-7’, **Fig. 1B, right**), RNA is extended to 6-mer and the 5’-end RNA base positions at fork loop 2 of the *β* subunit and the downstream edge of the transcription bubble. The 3’ end of the RNA is in a post-translocated state, forming a base pair with +3C tDNA, while +4G tDNA positions at the *i*+1 site as observed in the RTC-5’. The NTP binding *i*+1 site is empty but traps pyrophosphate. The +4G tDNA is waiting for GTP for further extension of the poly-G transcript. We also prepared the RTC by soaking GTP for 2 hours, but there was no further RNA extension beyond 8-mer RNA (data not shown) as observed in the RTC-30’, indicating that the 8-mer RNA is the longest RNA produced by *in crystallo* transcription. A series of structures of RTC show that 1) extra G bases are added at the RNA 3’ end by the G-G mismatch between the +4G tDNA and incoming GTP; and 2) the fitting of −4G RNA in the RIF-binding pocket is an obligatory step before RNA extension beyond 4-mer RNA.

### Comparison of the distances between bases to characterize hydrogen bonds

We captured the progression of RNA synthesis, from 2-mer to 8-mer, within the RTC crystals. In case of transcription from the *pyrG* promoter, the first three bases of RNA are synthesized by canonical transcription (*e.g.* RTC-3’) and then RNA synthesis is continued by reiterative transcription (*e.g.* RTC-5’, RTC-7’ and RTC-30’) (**Fig. 1B**). To gain insight into the RNA slippage mechanism, we analyzed the DNA and RNA base pairing at the *i* site of these structures. We assessed the distance between bases and the planarity of the base pair, which could be affected by GTP binding at the *i*+1 site during canonical and reiterative transcription. The distance between N1 of the G base of the RNA and N3 of the C base of the tDNA accommodated at the *i* site of RTC-3’ is 3.0 Å whereas, it is extended to 3.4 Å in case of the RTC-5’ containing the G-G mismatch at the *i*+1 site (**Fig. 1C, left and middle**). An average distance between C (DNA) and G (RNA) bases in atomic resolution DNA/RNA hybrid crystal structures is 2.9 Å with a minimum 2.75 Å and maximum 3.15 Å (19–21). We therefore concluded that the base pair between tDNA (+3C) and 3’ end of RNA is wobbled when RNAP switches the mode of RNA synthesis from canonical to reiterative transcription. The distance between C-G bases at the *i* site returns to 2.9 Å without GTP bound at the active site (RTC-7’, **Fig. 1C, right**) or when the RTC has completed poly-G RNA extension (RTC-30’) (17). Not only a distance between bases, but also planarity of base pair at the *i* site, is impaired during reiterative transcription. In the RTC-5’, +3C base of the tDNA is tilted about 15° toward the incoming GTP bound at the *i*+1 site (**Fig. 1C, middle**). A major difference found in the RTC-5’ compared with other RTC structures is the presence of an incoming GTP at the *i*+1 site, forming a G-G mismatch with the tDNA +3G base, which may wobble base pairing at the *i* site and initiate RNA slippage.

### Monitoring DNA translocation state during reiterative transcription in solution using 2-aminopurine fluorescence signal

The RTC-5’, RTC-7’ and RTC-30’ structures show that the +4G DNA base is translocated into the *i*+1 site when extra G residues are added to RNA during reiterative transcription (**Fig. 1B**). To validate this observation in solution, we monitored tDNA position during transcription by fluorescence signal of 2-aminopurine (2-AP) incorporated in the tDNA. 2-AP is an adenine analog (**Fig. 2A**) (22) and its fluorescence intensity is affected by its environment (23, 24). 2-AP displays weak and strong fluorescence signals when it stacked and unstacked with neighboring bases, respectively (**Fig. 2B and S3**) (25, 26).

**Fig. 2.**
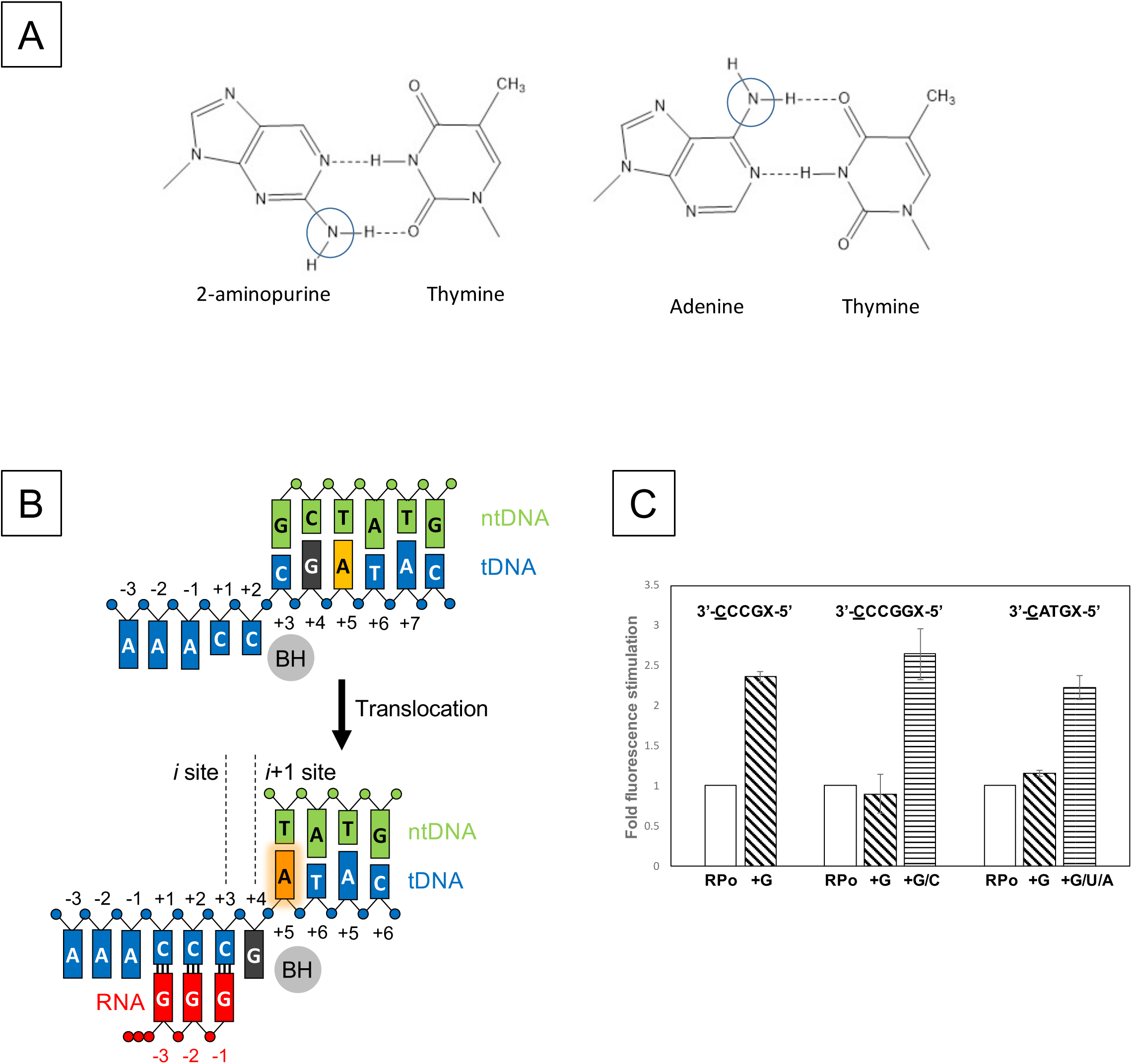
Monitoring equilibrium DNA translocation state in solution using 2-Aminopurine. **A)** The fluorophore 2-AP mimics the structure of natural adenine base and participates in the Watson-Crick interaction with thymine. **B)** Reaction scheme of reiterative transcription at the *pyrG* promoter. **C)** Fluorescence signal from 2-AP substituted at +5 or +6 position of the template strand DNA. Data are shown as mean ± SEM.

We prepared two DNA scaffolds containing a guanine DNA base as a quencher of 2-AP fluorescence with 2-AP at the +4 and +5 positions of the tDNA, respectively (**Table S2**). One scaffold contains the tDNA sequence 3’-<u>C</u>ATGX-5’ (transcription start site is underlined, X=2-AP) for canonical transcription (**Fig. S3B**) and another scaffold contains a RNA slippage prone tDNA sequence, 3’-<u>C</u>CCGX-5’, for reiterative transcription (**Fig. 2B**). For this assay, we used the *E. coli* RNAP σ^70^ holoenzyme since it is proficient at reiterative transcription from the *pyrG* promoter (17).

In the case of the canonical transcription DNA template, 2-AP displays low fluorescence in the RNAP-DNA complex (2-AP positioned at +5 site remains stacked with a quencher G base at +4 site) whereas it shows increased fluorescence after adding GTP, UTP and ATP in the RNAP-DNA complex (**Fig. 2C**). The result indicates that after 3-mer RNA synthesis, the +4G DNA base moves to the active site of RNAP (*i+1* site), leaving the 2-AP unstacked with +4G (**Fig. S3B**).

In the case of the reiterative transcription DNA template, 2-AP fluorescence increases upon addition of GTP to the RNAP-DNA complex for synthesizing 3-mer or longer RNA (**Fig. 2C**), demonstrating that the +4G tDNA is translocated to the *i*+1 site of the RNAP active site, which is consistent with the observation from the structural analysis of RTC (**Fig. 1**).

We also tested another DNA scaffold containing the *pyrG* transcription start site sequence but with an extra G base after +4G tDNA (3’-<u>C</u>CCGGX-5’, X = 2-AP) as a control, which would maintain the base stacking interaction between +5G and 2-AP while RNAP is engaged in reiterative transcription (**Fig. S3A**). When only GTP is mixed to the RNAP-DNA complex, 2-AP displays low fluorescence while the solution containing GTP and CTP that allows for translocation of +5G and 2-AP at the *i*+1 and *i*+2 sites, respectively, shows increased fluoresce signal (**Fig. 2C**).

### Kinetics of transcription-induced increase in fluorescence of 2-AP promoter DNA

A series of the RTC structures determined in this study revealed that RNAP requires 4∼5 min to synthesize 5-mer RNA from the *pyrG* promoter by *in crystallo* transcription, which is substantially slower than RNA synthesis from a promoter without a slippage prone sequence. For example, the structure of the initially transcribing complex containing 6-mer RNA (PDB: 4Q5S) (27), which contains the tDNA sequence 3’-<u>T</u>GAGTGC-5’, requires only 20 sec to produce 6-mer RNA *in crystallo* in the presence of ATP, CTP and UTP.

To measure the speed of RNA synthesis from slippage prone DNA in solution, we monitored the fluorescence of 2-AP embedded in tDNA at the +5 position with stopped-flow technique. We used the same DNA (**Table. S2**) for determining the DNA translocation state during reiterative and non-reiterative transcription. These DNA templates require at least 3-mer RNA synthesis for enhancing the 2-AP fluorescence signal. The data were best fit to a single exponential equation and kinetic values of the fluorescence signal from 2-AP are shown in **Fig. 3**. The rates of DNA translation of the RNA slippage prone and the canonical transcription DNA templates are ∼0.401 s^-1^ and ∼1.00 s^-1^, respectively, indicating that RNA synthesis from the slippage prone DNA is substantially slower than from the DNA for canonical transcription (**Fig. 3**). We also observed that the 2-AP fluorescence signal remains plateau, indicating that there is no backtracking of DNA:RNA hybrid during the reiterative transcription.

**Fig. 3.**
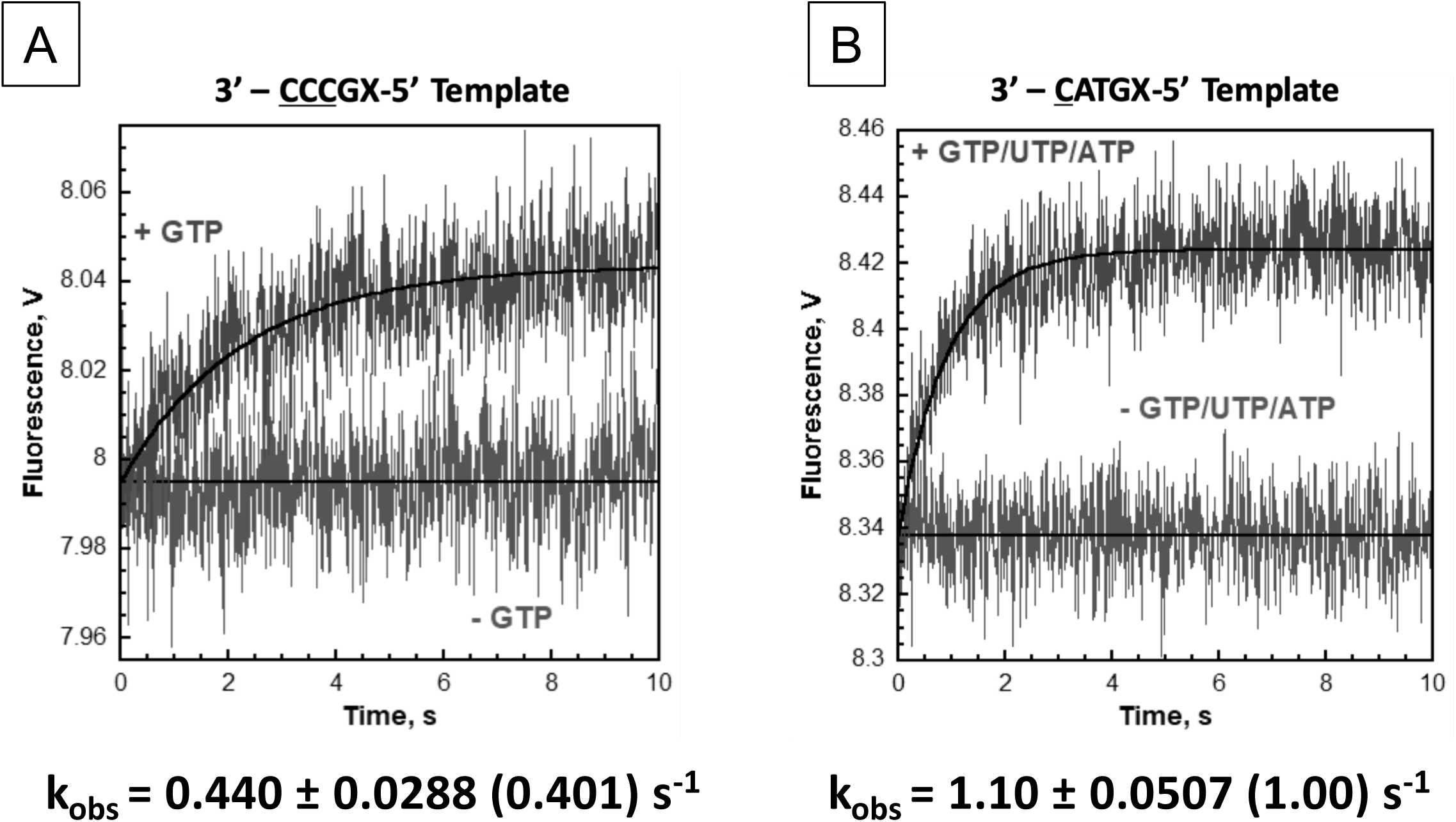
Kinetics of change in 2-AP fluorescence as measured by stopped-flow analysis. Time dependence of the increase in fluorescence upon mixing nucleotides to the reiterative transcription DNA template (A) and the canonical transcription DNA template (B). Each fluorescence trace represents the average of at least seven shots. For each DNA template, the observed increase in fluorescence was fit to a single exponential rise. If NTPs were omitted, a change in the fluorescence signal was not observed. The kinetic values are indicated.

### A role of the upstream sequences of the transcription start site of *pyrG* promoter

Reiterative transcription from the *pyrG* promoter places the −4G RNA base in the RIF-binding pocket. The RTC-7’ and RTC-30’ structures showed the −1A base of tDNA partially overlapping with the −3G base of RNA (**Fig. 4A**). Such base stacking is only possible in the presence of a purine (tDNA) and purine (RNA) combination (adenine in tDNA and guanine in RNA in the case of *pyrG* promoter transcription) (**Fig. 4B**). Consistent with this fact, the upstream sequence of the transcription start site of the *pyrG* promoter is highly conserved in other closely related bacteria, and particularly, tDNA bases at positions −1 and −2 are adenine in a majority of promoters (**Fig. S4**). We therefore hypothesized that the −1A tDNA may block RNA extension toward the RNA exit channel, thereby the −4G RNA base is pushed into the RIF-binding pocket when RNA slips on the tDNA.

**Fig. 4.**
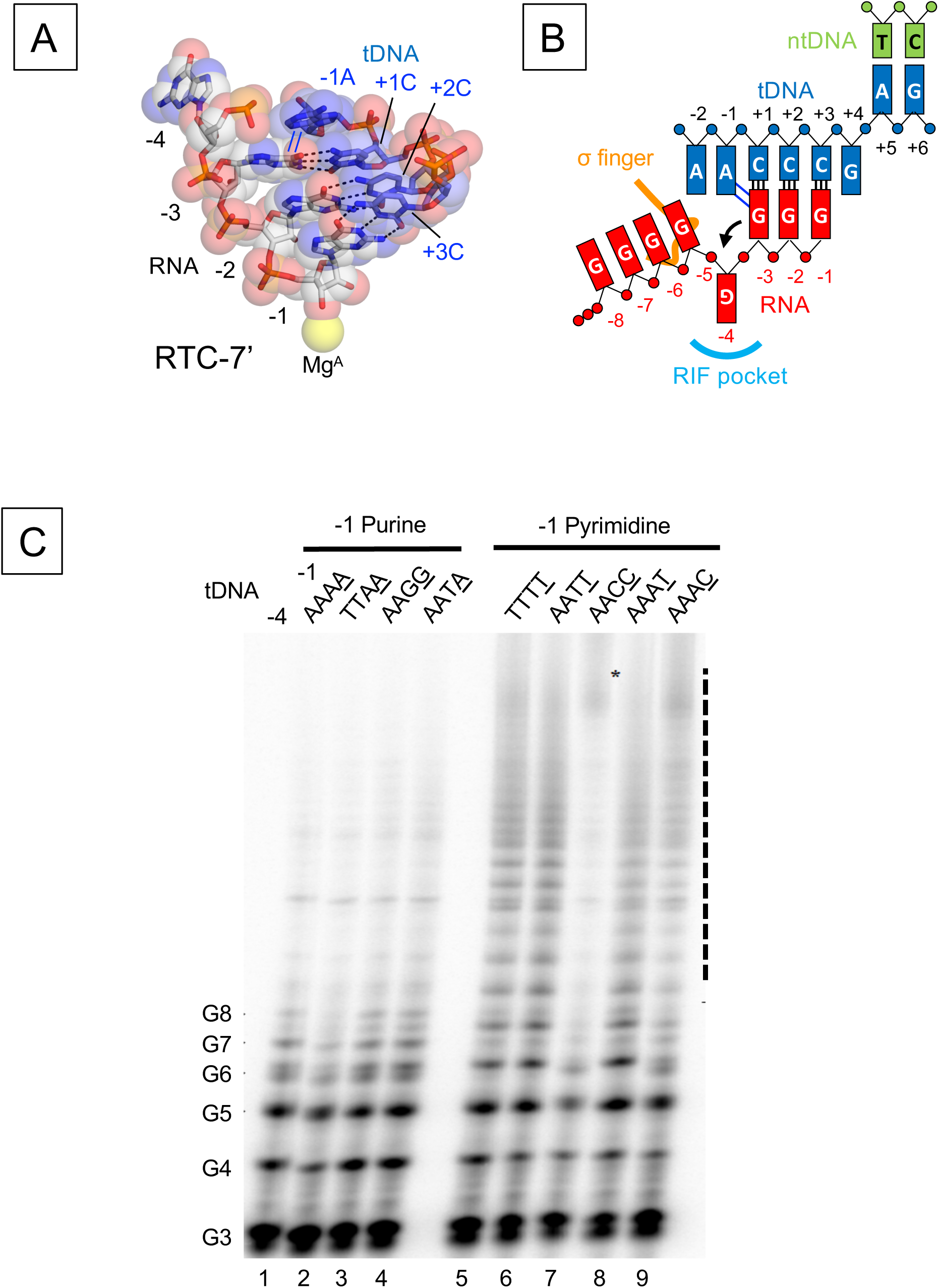
Role of the −1 template DNA base in reiterative transcription **A)** DNA and RNA hybrid region of the RTC-7’ structure. Base parings and base stacking interactions are depicted as black dashed lines and blue lines, respectively. **B)** Schematic representation of reiterative transcription from the *pyrG* promoter and the base stacking interaction between the −1 tDNA adenine base and the guanine base of RNA at −1 position. **C)** *In vitro* transcription assays measuring poly-G RNA production at the *pyrG* variant promoters. Sequences of tDNA from −4 to −1 positions used for the transcription assay are indicated above each lane. ^32^p-labelled GpGpG primer (20 µM), corresponding to positions +1 to +3 of the promoter in the presence of GTP (100 µM) was used. Positions of the poly-G products are indicated. Poly-G RNAs longer than 8 bases are indicated as a dashed line.

First, we investigated a role of adenine bases of tDNA by *in vitro* transcription (**Fig. 4C**). The *pyrG* promoter from *B. subtilis* contains 4 adenine bases from the −4 to −1 positions in the tDNA. We prepared a series of *pyrG* promoter variants with substitutions from the −4 to −1 positions (**Table. S3**) and tested their abilities to produce reiterative transcripts. The reaction was performed in the presence of a GpGpG trinucleotide primer complementary to the tDNA positions from +1 to +3 and GTP for efficient RNA extension. The wild-type promoter produces poly-G RNA around 8 bases in length, and adenine bases at −4, −3 and −2 tDNA can be substituted with thymine without changing the activity of reiterative transcription (**Fig. 4C, lanes 1, 2 and 4**). Guanine substitutions at −2 and −1 positions of tDNA did not influence the transcription (**Fig. 4C, lane 3**). In contrast, *pyrG* derivatives containing thymine or cytosine substitutions at the −1 position increased the length of poly-G RNA transcript substantially (20-mer or longer) (**Fig. 4C, lanes 5-9**). These results indicate that the tDNA base at the −1 position determines the length of reiterative RNA product from the *pyrG* promoter; short RNAs are synthesized in the presence of adenine and guanine (purine bases) and longer RNA are produced in the presence of thymine and cytosine (pyrimidine bases).

### Structural analysis of the role of −1 base of template DNA during reiterative transcription

To understand how the −1 base of tDNA determines the length of reiterative transcription products, we solved the crystal structures of RTC containing *pyrG* variants having tDNA bases of −1G, −1C and −1T (**Fig. 5A**, and **Table S5**). All RTC structures show electron densities corresponding to poly-G RNA products starting from the RNAP active site; however, the position of the 5’ end of RNA is different depending on the sequence of tDNA. In the case of the RTC containing −1G tDNA (RTC-1G) (**Fig. 5A, left)**, the RNA is accommodated as found in the wild-type RTC; the RNA forms a 3 bp hybrid with tDNA, the −3G RNA base partially overlaps with - 1G tDNA base, the −4G RNA base fits into the RIF-binding pocket, and the 5’ end of RNA extends toward the main channel (**Fig. 5B, left**). In sharp contrast, the RTCs with −1C and −1T tDNA bases (RTC-1C and RTC-1T) (**Fig. 5A, middle and right**) contain a 4 bp DNA/RNA hybrid; −4G RNA base forms a Watson-Crick base pair (C-G) with tDNA base (−1C) in the RTC-1C and a wobble base pair (T-G) with tDNA base (−1T) in the RTC-1T (**Fig. 5B, middle and right**). In both structures, the triphosphate of RNA contacts a tip of σ finger (residues 321-327). The path of RNA in the RTC-1C and RTC-1T is the same as the initially transcribing complex containing 6 bp DNA/RNA hybrid (PDB: 4G7H) (27).

**Fig. 5.**
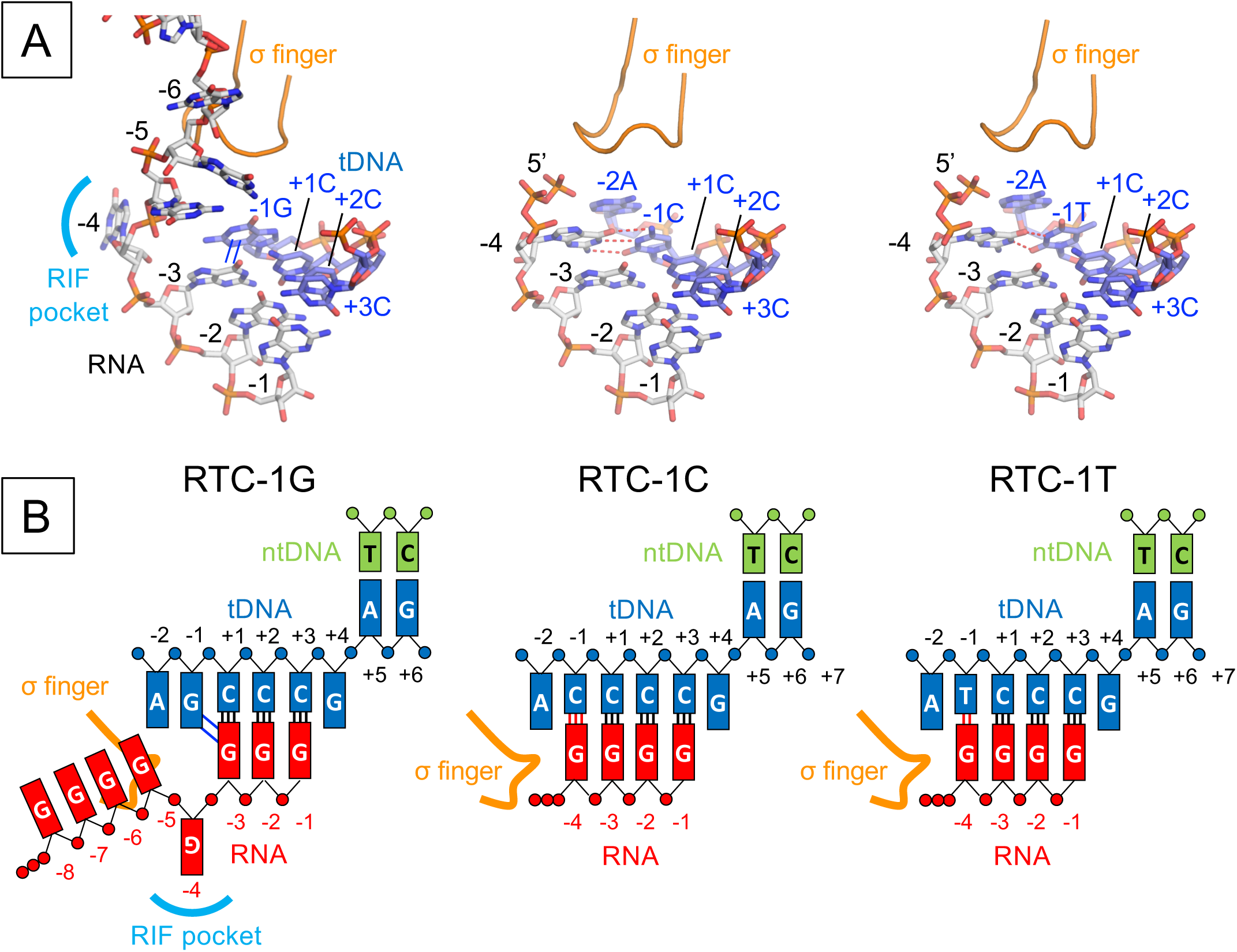
The structures of reiterative transcription complex from the *pyrG* promoter variants. A) Structures of the RTC-1G (left), RTC-1C (middle) and RTC-1T (right). RNA and tDNA are shown as stick models and the σ finger is depicted as a ribbon model. Base stacking and base paring interactions are depicted as blue lines (in RTC-1G) and red dashed lines (in RTC-1C and RTC-1T), respectively. B) Schematic representations of reiterative transcription from the *pyrG* promoter variants containing −1G (RTC-1G, left), −1C (RTC-1C, middle) and −1T (RTC-1T, right).

### Structure of a reiterative transcription complex at the *pyrBI* promoter

Reiterative transcription from the *pyrBI* operon of *E. coli* produces much longer RNA products in the presence of ATP and high concentration of UTP (**Fig. S5B**) (16), which is akin to the transcription from the *pyrG* promoter variant containing a pyrimidine base at −1 tDNA (**Fig. 4C**). To study similarity and difference of reiterative transcription from the *pyrBI* and *pyrG* promoters, we determined the crystal structures of the open complex containing the *pyrBI* promoter and its RTC by soaking an open complex crystal into a solution containing ATP and UTP (**Fig. 6 and Table S6**). In the open complex structure, thymine bases of tDNA position at the *i* and *i*+1 sites of the RNAP active site, showing that the RNA synthesis starts from +1 position. The RTC structure shows the RNAP active site containing 4-mer RNA (5’-AAUU) with an UTP positioned at the pre-insertion site (**Fig. 6B**). RNA extends directly toward the *σ* finger and steric clash between the RNA triphosphate and *σ* finger prohibits RNA extension beyond 4-mer RNA.

**Fig. 6.**
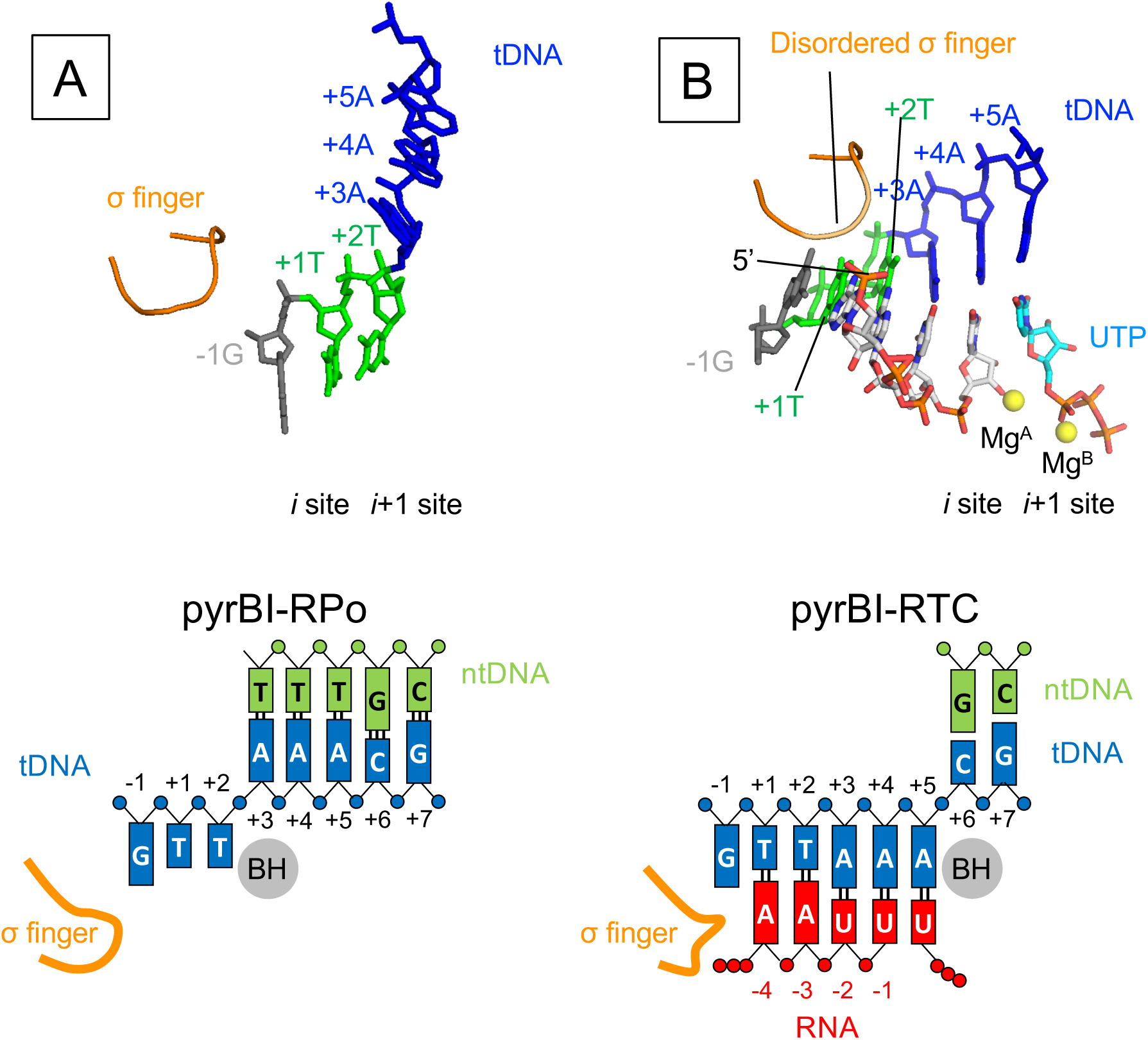
The structures of the reiterative transcription complex from the *pyrBI* promoter. Structures of the RNAP-pyrBI promoter binary open complex (pyrBI-RPO, **A**) and reiterative transcription complex (pyrBI-RTC, **B**). RNA and tDNA are shown as stick models and the σ finger is depicted as a ribbon model. The Mg ions bound at the active site of RNAP are shown as yellow spheres. Schematic representations of the pyrBI-RPO (left) and pyrBI-RTC (right) are shown at lower panels.

## DISCUSSION

### Structural and biochemical evidence of base sharing during RNA slippage

In this study, using time-dependent soak-trigger-freeze X-ray crystallography, we analyzed how RNAP synthesizes RNA from the *pyrG* promoter. The first three RNA bases are synthesized by the canonical transcription, which is a simple copy of DNA sequence to RNA. In the case of a CTP limited condition, RNAP extends RNA with GTP by reiterative transcription (**Fig. S1**). The structure of RTC-5’ containing 4-mer RNA, which represents the transcription complex right after the mode of RNA synthesis has switched, revealed that +4G tDNA is positioned at the active site of RNAP (*i*+1 site) and forms a mismatch pair with an incoming GTP using an *anti-anti* conformation (**Fig. 1B, middle**). The structure also revealed that a base pair between tDNA (+3C tDNA) and the 3’ end of RNA positioned at the *i* site is wobbled (N3 of C and N1 of G, 3.4 Å) and the +3C tDNA base is tilted toward an incoming GTP, likely due to a mismatch between +4G tDNA and incoming GTP (**Fig. 1C, middle**). The opening of the trigger loop causes the *α*-phosphate of GTP to be 6.1 Å away from the 3’-OH of RNA and maintain GTP in a nonreactive state for the nucleotidyl transfer reaction. We speculate that the trigger loop would then adapt the completely closed conformation, forming the trigger helix, to load the GTP in the reactive state followed by the nucleotidyl transfer reaction (**Fig. 7, top**). RNA extension may trigger base sharing of the +3C tDNA with guanine bases of RNA at the *i* and *i*+1 sites and would promote shifting of the DNA/RNA hybrid in a stepwise manner (**Fig. 7, middle**). This hypothesis is supported by the observation that the base pairs upstream from the *i* site are not wobbled. After RNA is extended, only RNA translocates in the upstream direction to prepare for the next cycle of reiterative transcription (**Fig. 7, bottom**). This prediction is supported by the results of 2-AP fluorescence based DNA translocation assay (**Figs. 2 and 3**). Extending RNA from 3-mer to 4-mer determines the fate of RNA synthesis from the *pyrG* promoter. Translocating +4G tDNA at the active site (*i*+1 site) after 3-mer RNA synthesis is also observed in the DNA translocation assay by monitoring the 2-AP fluorescence signal from tDNA (**Fig. 2**). The +4G tDNA moving at the active site is essential for regulating *pyrG* expression depending on CTP availability; when the CTP concentration is high enough, CTP is loaded at the active site, forms a base pair with +4G tDNA and RNAP undergoes canonical transcription (**Fig. S1**). Previous study by Meng et al observed the similar reiterative transcription pattern when the +4G tDNA was replaced with either A or T (6), further suggesting that the mismatch between the +4 tDNA and incoming GTP favors transcript slippage at *pyrG* promoter.

**Fig. 7.**
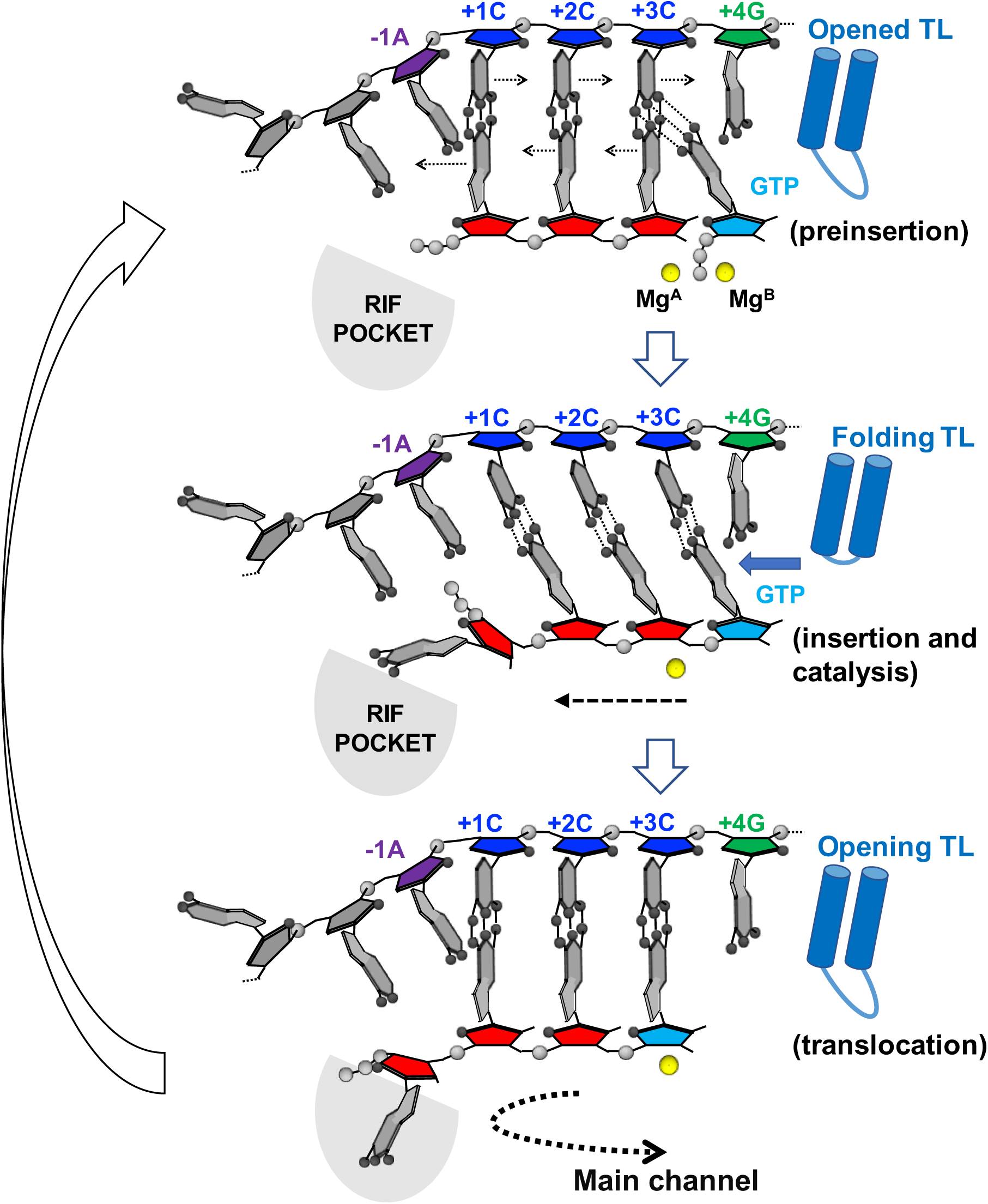
Proposed model of RNA slippage during reiterative transcription from the *pyrG* promoter. Cartoon model of the active site of RNAP. *pyrG* initially transcribed region sequence (template DNA, −1A: purple, +1C, +2C, +3C: blue, +4G: green), nascent RNA (5’-GGG-3’, red), incoming nucleotide (GTP, cyan), active site Mg^2+^ (yellow sphere), trigger loop (blue shape), and rifampin-binding pocket (RIF POCKET, gray) are shown.

### The nature of the base at −1 position of template DNA determines initial pathway of poly-G transcript

*In vitro* transcription assays (**Fig. 4C**) and the structures of the RTCs containing *pyrG* promoter variants (**Fig. 5**) revealed the nature of tDNA base at the −1 position, either purine or pyrimidine, determines the fates of poly-G transcripts regarding length and direction. In the case of tDNA having a purine base (guanine or adenine) at the −1 position, RNAP synthesizes RNA around 8-mer in solution (**Fig. 4C, lanes 1-4**) and *in crystallo*. The guanine base at the 5’ end of the RNA sterically clashes with a purine base of tDNA at −1 position when RNA slips on the tDNA for the first time, resulting in flipping an RNA base into the RIF-binding pocket (**Figs. 4A and B**). It is important to note that the steric collision between the RNA and tDNA bases only happens during reiterative transcription. Further reiterative transcription extends RNA toward the main channel of RNAP until the poly-G RNA eventually switches to canonical transcription when a CTP molecule is incorporated into the 3’ end of the transcript (**Fig. S1**).

In the case of tDNA having a pyrimidine base at the −1 position (−1C and −1T), RNAP synthesizes RNA over 40-mer in solution (**Fig. 4C, lanes 5-8**) but only 4-mer *in crystallo* (**Fig. 5**). The structures show that a pyrimidine base of tDNA does not hinder RNA movement toward the dedicated RNA exit channel, which allows for a base pair between the 5’ end of RNA and the −1 tDNA base right after the first RNA slippage event (Watson-Crick or wobble base pair in case of −1C and −1T, respectively), resulting in forming 4 bp DNA/RNA hybrid (**Fig. 5**). The reiterative transcript extends toward the RNA exit channel and the 5’ end of RNA clashes with the σ finger, which may trigger the release of σ factor from RNAP core enzyme and allow for longer RNA extension. RNAP can extend RNA only 4-mer *in crystallo* because σ cannot be released from the core enzyme due to crystal packing.

It has been demonstrated that two short segments of the 5’ untranslated region (UTR) of *pyrG* are required for CTP-dependent regulation, including the initially transcribed region (ITR) of the *pyrG* promoter (5’-<u>G</u>GGC, transcription start site is underlined) and the pyrimidine rich sequence of the attenuator (**Fig. S1**) (6, 28). In addition to these DNA cis elements, in this study, we shed light on the function of the purine rich sequence of tDNA just upstream of the transcription start site. These bases, particularly a purine base at −1 tDNA position, maintain the length of DNA/RNA hybrid to 3 bases and guide the 5’-end of RNA toward the RIF binding pocket of RNAP, which is an obligatory step for RNA extension toward the main channel of RNAP. Reiterative RNA synthesis pauses around 8∼10 bases when the 5’-end RNA reaches the narrow opening of the main channel of RNAP, providing time for CTP to be incorporated at the 3’ end of RNA and for RNAP to switch the mode of RNA synthesis from the reiterative to canonical transcription. Maintaining the 3 base DNA/RNA hybrid length in the *pyrG* reiterative transcription complex is critical to the regulation of *pyrG* gene expression depending on the CTP availability. In the ITR, not only a run of three but also a run of four or five G residues permits reiterative transcription; however, these extra Gs in the ITR result in less than optimum regulation of *pyrG* expression under CTP limited conditions. A run of five or more G residues suppresses RNA slippage due to suppressing the DNA/RNA hybrid melting (28).

### Reiterative transcript initiation from other promoters

The mechanism of RNA extension observed for the *pyrG* promoter will not hold for other promoters engaged in reiterative transcription, such as *pyrBI* (5’-<u>A</u>A*TTT*G: transcription start site is underlined and slippage prone sequence is italicized) (16), *codBA* (5’-<u>A</u>*TTTTTT*G) (29), and *upp-uraA* (5’-<u>G</u>A*TTTTTTTT*G) (30) in *E. coli*, because these promoters produce 5∼10 bases of RNA before slipping and synthesize much longer stretches of reiterative RNAs (30 nucleotides or longer) (**Fig. S5**). Furthermore, once reiterative transcription starts from these promoters, RNA synthesis does not switch to canonical transcription (**Fig. S2**). The mechanism of reiterative transcription from these promoters might be similar to *pyrG* promoter variants containing −1T or - 1C as investigated in this study (**Figs. 4 and 5**); these variants produce much longer reiterative RNA relative to the wild-type *pyrG* promoter. This hypothesis is consistent with our structural observation that the crystal containing the *pyrBI* promoter in the presence of ATP and UTP substrates formed 4-mer RNA (5’-AAUU-3’) and its 5’ end collides with the *σ* finger (**Fig. 6**). Investigating how the *pyrBI*-type RTC accommodates a long stretch of reiterative RNA cannot be addressed by *in crystallo* transcription and X-ray crystallography structure determination because *σ* release is not permitted in the RNAP crystals. Elucidating the structural basis of *pyrBI*-type reiterative transcription could be achieved by determining the structure of RTC by cryo-electron microscopy (cryo-EM), a powerful method to determine high resolution macromolecular structures in solution (31).

## Experimental procedures

### Preparation and Purification of *T. thermophilus* and *E. coli* RNAPs

*T. thermophilus* and *E. coli* RNAP holoenzymes were prepared as described previously (27, 32).

### Preparation of promoter DNA scaffolds for the crystallization, the DNA translocation assay and the *in vitro* transcription assay

The promoter DNA scaffold that resembles the *B. subtilis pyrG* promoter region and its variants, and the *E. coli pyrBI* promoter region were constructed using two oligodeoxynucleotides for template and non-template DNA strands. The DNA oligonucleotides used for the crystallization, the DNA translocation assay, and the *in vitro* transcription assay are shown in the **Table S1, S2, S3**, respectively. DNA strands were annealed in 40 μL containing 10 mM Tris-Hcl (pH 8.0), 50 mM NaCl, and 1 mM EDTA to the final concentration of 0.5 mM. The solutions were heated at 95 °C for 10 min and then the temperature was gradually decreased to 22 °C.

### Crystallization of the *T. thermophilus* RNAP promoter DNA complexes

The crystals of the RNAP and promoter DNA complex were prepared as described previously (17). To prepare the crystals of RTC from the *pyrG* promoter, the RNAP and *pyrG* DNA complex crystals were transferred to cryoprotection solution containing 1 mM GTP, harvested from the soaking solution at indicated time points, and flash frozen in liquid nitrogen. To prepare the crystals of RTC from the *pyrBI* promoter, the RNAP and *pyrBI* DNA complex crystal was transferred to a cryoprotection solution containing 5 mM ATP, 5 mM UTP and 500 μM GTP for 1 hour and then frozen by liquid nitrogen.

### X-ray data collections and structure determinations

The X-ray datasets were collected at the Macromolecular Diffraction at the Cornell High Energy Synchrotron Source (MacCHESS) F1 beamline (Cornell University, Ithaca, NY) and structures were determined as previously described (17, 27) using the following crystallographic software: HKL2000 (33), Phenix (34) and Coot (35).

### *In vitro* transcription assay

The transcription assays on the *pyrG* promoter and its variants were performed in 10 μL containing 250 nM RNAP holoenzyme, 250 nM DNA, 100 μM GTP and ^32^P-labelled GpGpG primer in the transcription buffer [40 mM Tris-HCl (pH 8 at 25 °C), 30 mM KCl, 10 mM MgCl2, 15 μM acetylated BSA, 1mM DTT]. RNAP-DNA-GpGpG primer were preincubated at R.T for 10min. After adding GTP, the samples were incubated at 37 °C for 10 mins and the reaction stopped by adding 10 μL of stop buffer (90% formamide, 50 mM EDTA, xylene cyanol and bromophenol blue). The transcription assay on the *pyrB*I promoter was performed in 10 μL containing 250 nM RNAP holoenzyme, 250 nM DNA, 100 μM GTP, 100 μM ATP, [γ-^32^p] ATP and 5 mM to 10 μM UTP. The reaction products were electrophoretically separated on a denaturing 24% polyacrylamide/7 M urea gel and visualized with a phosphorimager (Typhoon 9410; GE Healthcare).

### Monitoring DNA translocation state during reiterative transcription in solution using 2-aminopurine (2-AP) fluorescence (equilibrium study)

Experiments were performed at 37 °C in the transcription buffer. *E. coli* RNAP holoenzyme (500 nM) and DNA (100 nM) was preincubated for 10 min at 25 °C for open complex formation. Transcription was initiated by mixing one or more indicated NTPs (each at 200 μM). After mixing, the fluorescence was detected at excitation and emission wavelengths of 315 nm and 375 nm, respectively, using Spectramax-M5 spectrophotometer (Molecular Devices).

### Measurement of DNA translocation kinetics during reiterative transcription

Stopped-flow studies were performed on an Applied Photophysics SX20 stopped-flow machine equipped with a fluorescence detector. All experiments were performed at room temperature (23 ± 2 °C) in the transcription buffer and the final ionic strength was adjusted to 100 mM by the addition of appropriate amounts of KCl. Syringe A (70 μL) containing the *E. coli* RNAP holoenzyme (500 nM) and DNA (100 nM) was mixed with an equal volume of Syringe B (70 μL) containing one or more NTPs (each at 200 μM). Upon mixing, 2-AP fluorescence was monitored by exciting at 315 nm and monitoring the emission using a 350 nm cutoff filter (Andover Corporation, Salem, NH). The fluorescence traces were recorded by collecting 1000 total time points over 10 s. All traces were analyzed using Applied Photophysics ProData^TM^ and Kaleidagraph softwares.

## AUTHOR CONTRIBUTIONS

Y.S., M.H. and K.S.M. contributed to the design of the experiments. Y.S. crystallized the RTCs and collected the X-ray diffraction data. Y.S. and K.S.M. determined X-ray crystal structures. Y.S. developed and conducted the *in vitro* transcription and 2-AP fluorescence assays. Y.S. and M.H. developed and conducted the stopped flow experiment. All authors participated in the interpretation of the results and in writing the manuscript.

## ACKNOWLEDGEMENTS

We thank the staff at the MacCHESS for support of crystallographic data collection. We thank Dr. Charles L. Turnbough, Jr, Catherine Sutherland and Shoko Murakami for critically reading the manuscript. This work was supported by NIH grants R01 GM087350 and R35 GM131860 (K.S.M.).

## SUPPORTING INFORMATION

**Table S1:**
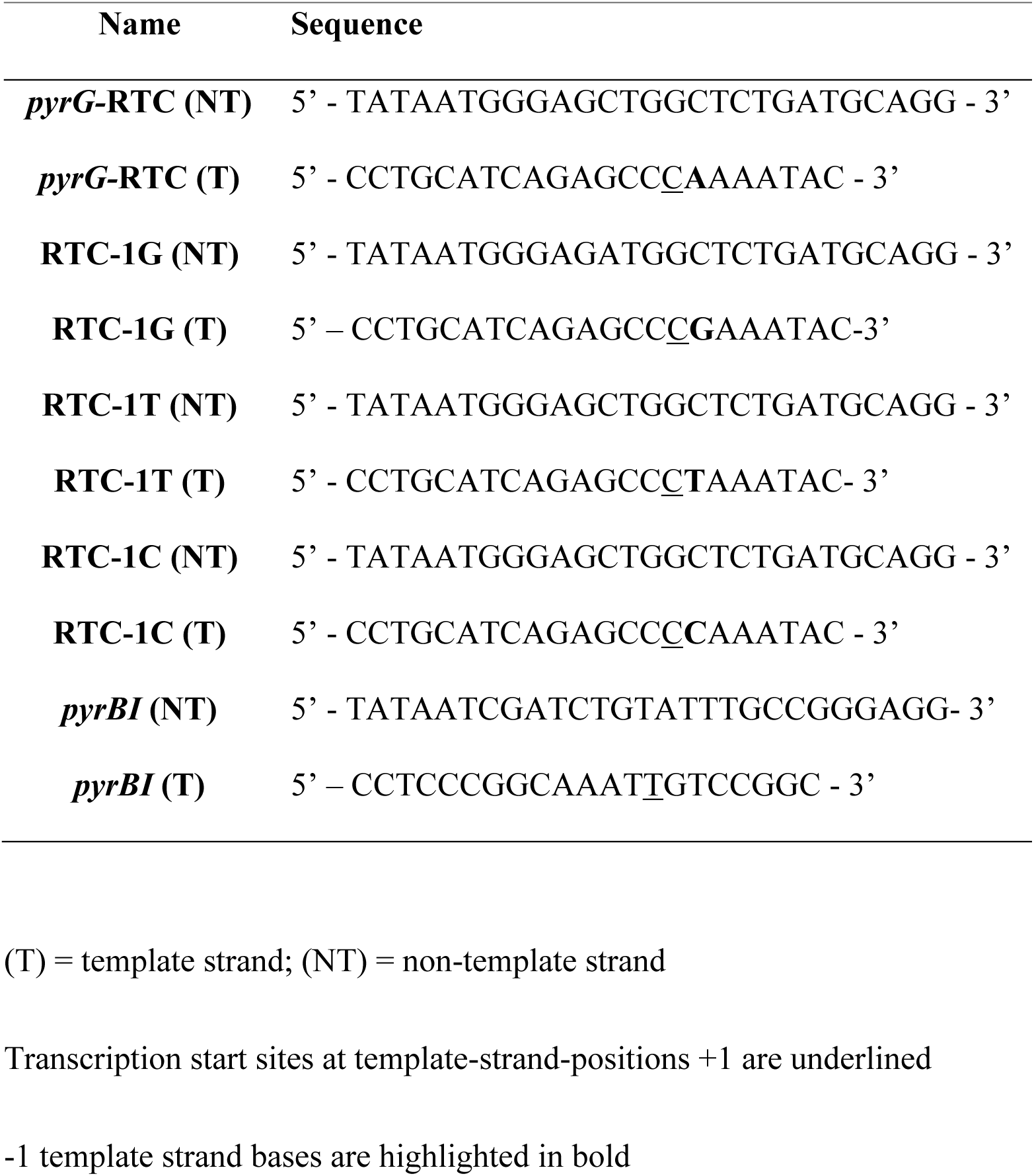
DNA oligonucleotides used for crystallization.

**Table S2:**
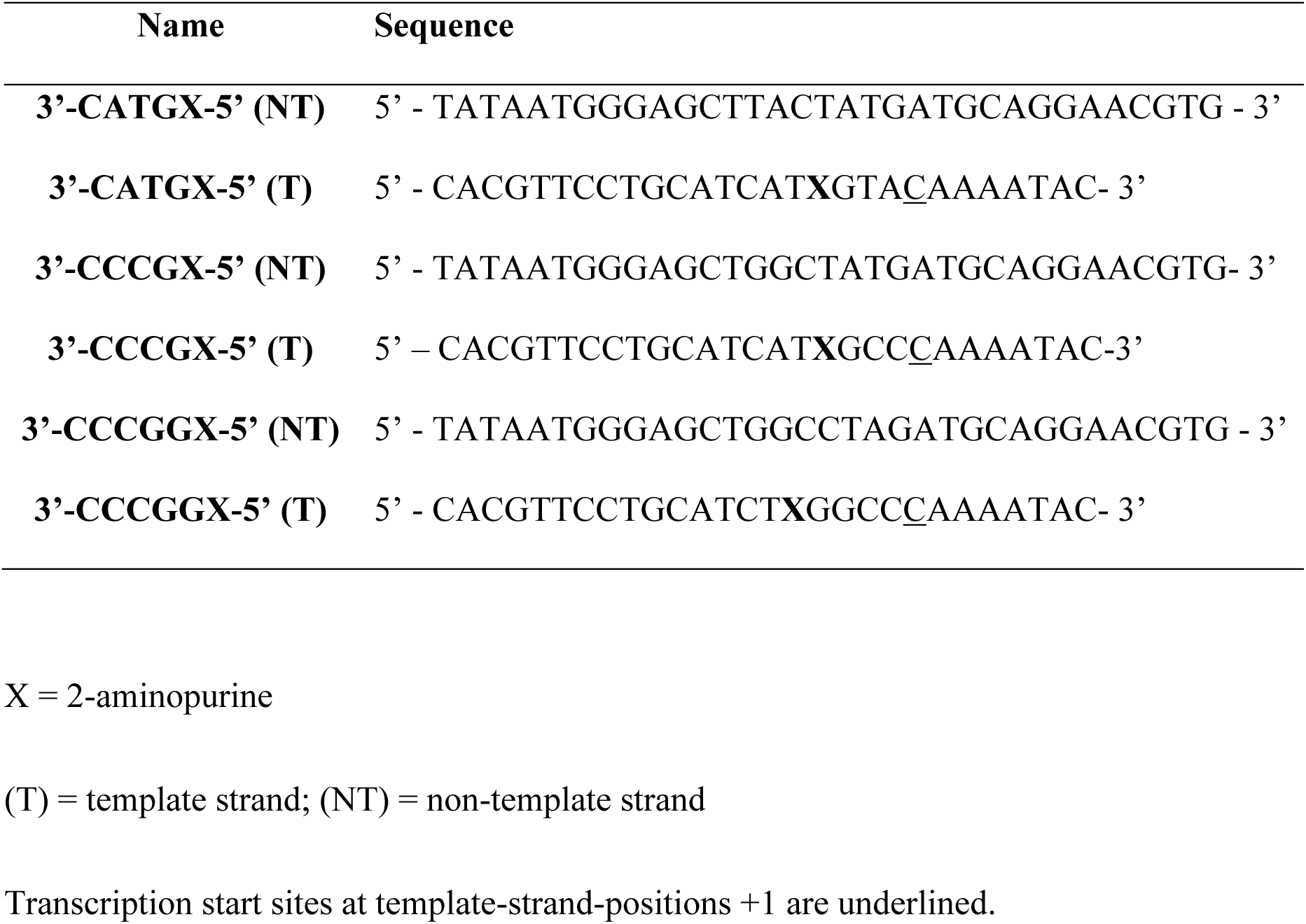
DNA oligonucleotides used for equilibrium and kinetics translocation assays using 2-aminopurine.

**Table S3:**
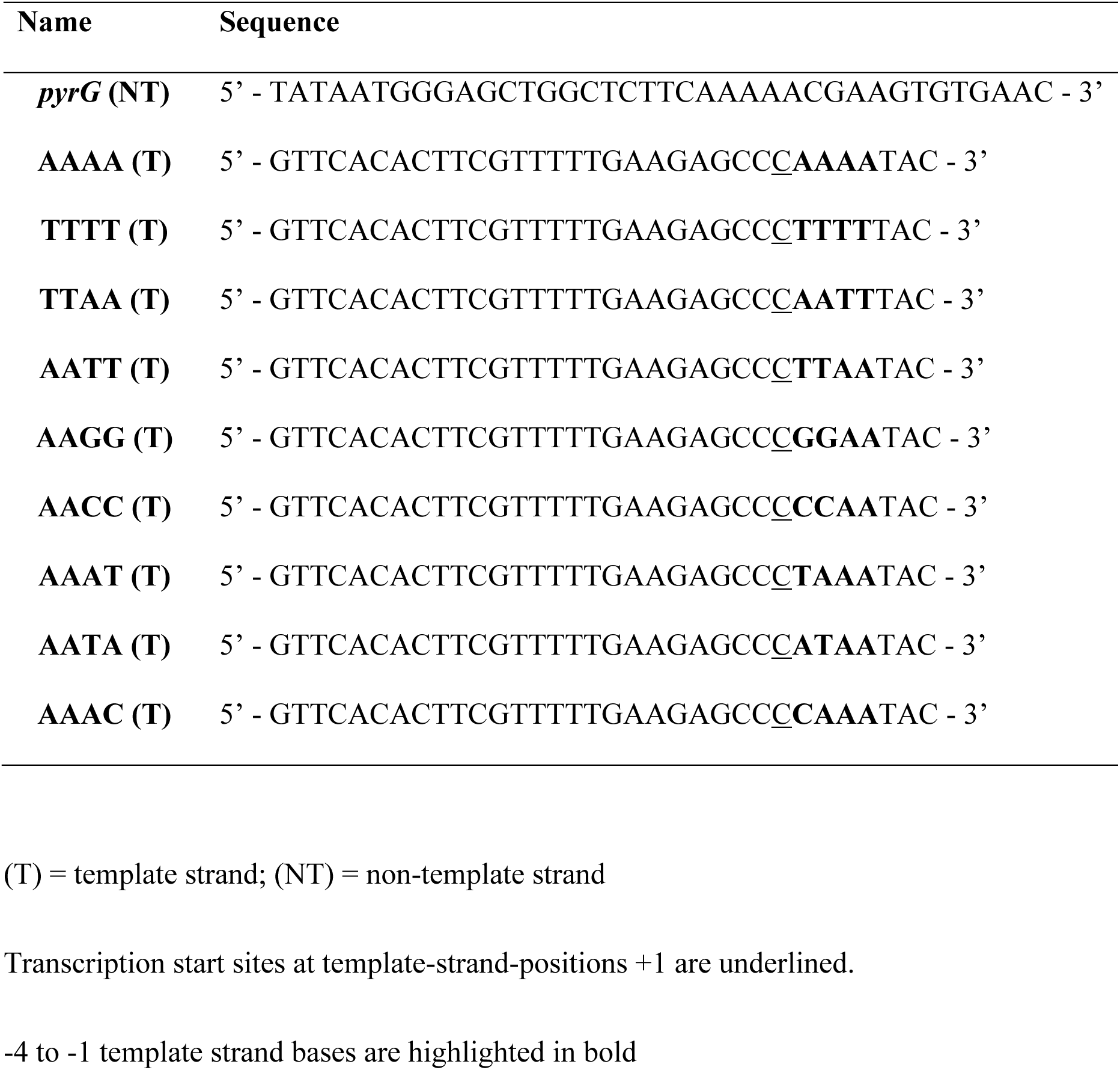
DNA oligonucleotides used for *in vitro* transcription assays.

**Table S4:** Data collection and refinement statistics of RTC intermediates.

| Complex<br>PDB code | RTC-3'<br>6OY5 | RTC-5'<br>6OY6 | RTC-7'<br>6OY7 |
| --- | --- | --- | --- |
| **Data collection** |  |  |  |
| Space group | C2 | C2 | C2 |
| Cell dimensions |  |  |  |
| *a* (Å) | 186.09 | 186.88 | 185.68 |
| *b* (Å) | 102.41 | 102.14 | 101.61 |
| *c* (Å) | 297.23 | 297.36 | 295.80 |
| $\beta$ (°) | 99.00 | 98.81 | 98.76 |
| Resolution (Å) | 50 – 3.0 | 50 – 3.1 | 50 – 3.04 |
| Total reflections | 295,628 | 261,015 | 316,859 |
| Unique reflections | 99,590 | 88,633 | 100,023 |
| Redundancy | 3.0 (2.5) | 2.9 (2.3) | 3.2 (2.7) |
| Completeness (%) | 90.0 (64.4) | 88.0 (56.0) | 95.1 (72.7) |
| *I* / $\sigma$ | 11.9 (0.85) | 9.64 (0.64) | 9.54 (0.91) |
| CC<sup>1/2</sup> | (0.544) | (0.488) | (0.520) |
| **Refinement** |  |  |  |
| Resolution (Å) | 44 – 3.1 | 44– 3.1 | 48 – 3.04 |
| *R*<sub>work</sub> | 0.207 | 0.212 | 0.207 |
| *R*<sub>free</sub> | 0.259 | 0.264 | 0.248 |
| No. of atoms | 28,534 | 28,578 | 28,534 |
| R.m.s deviations |  |  |  |
| Bond length (Å) | 0.011 | 0.012 | 0.012 |
| Bond angles (°) | 1.546 | 1.551 | 1.612 |
| Clashscore | 10.6 | 9.39 | 12.22 |
| Ramachandran favored, % | 95.96 | 95.96 | 95.45 |
| Ramachandran outliers, % | 0.78 | 0.75 | 1.10 |

Data sets were collected at MacCHESS F1 line, Ithaca, NY
\*Highest resolution shells are shown in parentheses

**Table S5:** Data collection and refinement statistics of −1 tDNA variants.

| Complex | -1G | -1C | -1T |
| --- | --- | --- | --- |
| PDB code | 6OVR | 6OVY | 6OW3 |
| **Data collection** |  |  |  |
| Space group | C2 | C2 | C2 |
| Cell dimensions |  |  |  |
| *a* (Å) | 185.90 | 184.83 | 186.19 |
| *b* (Å) | 101.42 | 102.48 | 101.83 |
| *c* (Å) | 295.20 | 295.60 | 295.76 |
| $\beta$ (°) | 98.64 | 98.80 | 98.64 |
| Resolution (Å) | 50 – 2.83 | 50 – 3.0 | 50 – 2.77 |
| Total reflections | 471,538 | 398,603 | 518,675 |
| Unique reflections | 123,691 | 106,213 | 136,486 |
| Redundancy | 3.8 (3.3) | 3.8 (3.0) | 3.8 (3.1) |
| Completeness (%) | 97.1 (76.6) | 97.6 (81.6) | 97.7 (78.2) |
| *I* / $\sigma$ | 16.3 (1.27) | 9.15 (1.21) | 16.24 (1.29) |
| CC<sup>1/2</sup> | (0.740) | (0.644) | (0.744) |
| **Refinement** |  |  |  |
| Resolution (Å) | 42 – 2.84 | 46 – 3.0 | 37 – 2.77 |
| *R*<sub>work</sub> | 0.221 | 0.213 | 0.220 |
| *R*<sub>free</sub> | 0.267 | 0.237 | 0.257 |
| No. of atoms | 28,535 | 28,273 | 28,430 |
| R.m.s deviations |  |  |  |
| Bond length (Å) | 0.013 | 0.012 | 0.012 |
| Bond angles (°) | 1.087 | 1.573 | 1.643 |
| Clashscore | 24.08 | 12.47 | 12.46 |
| Ramachandran favored, % | 93.9 | 92.9 | 92.5 |
| Ramachandran outliers, % | 0.87 | 0.9 | 1.1 |

Data sets were collected at MacCHESS F1 line, Ithaca, NY
\*Highest resolution shells are shown in parentheses

**Table S6:** Data collection and refinement statistics of *pyrBI*.

| Complex<br>PDB code | *pyrBI*-RPo<br>6P70 | *pyrBI*-1h<br>6P71 |
| --- | --- | --- |
| **Data collection** |  |  |
| Space group | C2 | C2 |
| Cell dimensions |  |  |
| *a* (Å) | 185.18 | 187.51 |
| *b* (Å) | 100.85 | 100.66 |
| *c* (Å) | 294.87 | 296.30 |
| $\beta$ (°) | 98.81 | 98.32 |
| Resolution (Å) | 50 – 3.05 | 50 – 2.85 |
| Total reflections | 342,091 | 435,858 |
| Unique reflections | 101,835 | 122,765 |
| Redundancy | 3.4 (2.9) | 3.6 (2.5) |
| Completeness (%) | 99.7 (97.0) | 96.2 (77.7) |
| *I* / $\sigma$ | 10.9 (1.21) | 9.86 (1.0) |
| CC<sup>1/2</sup> | (0.730) | (0.647) |
| **Refinement** |  |  |
| Resolution (Å) | 46 – 3.05 | 49 – 2.92 |
| *R*<sub>work</sub> | 0.207 | 0.211 |
| *R*<sub>free</sub> | 0.253 | 0.249 |
| No. of atoms | 28,359 | 28,417 |
| R.m.s deviations |  |  |
| Bond length (Å) | 0.011 | 0.012 |
| Bond angles (°) | 1.451 | 1.513 |
| Clashscore | 10.14 | 6.83 |
| Ramachandran favored, % | 97.1 | 97.2 |
| Ramachandran outliers, % | 0.32 | 0.37 |

Data sets were collected at MacCHESS F1 line, Ithaca, NY
\*Highest resolution shells are shown in parentheses

**Fig. S1.**
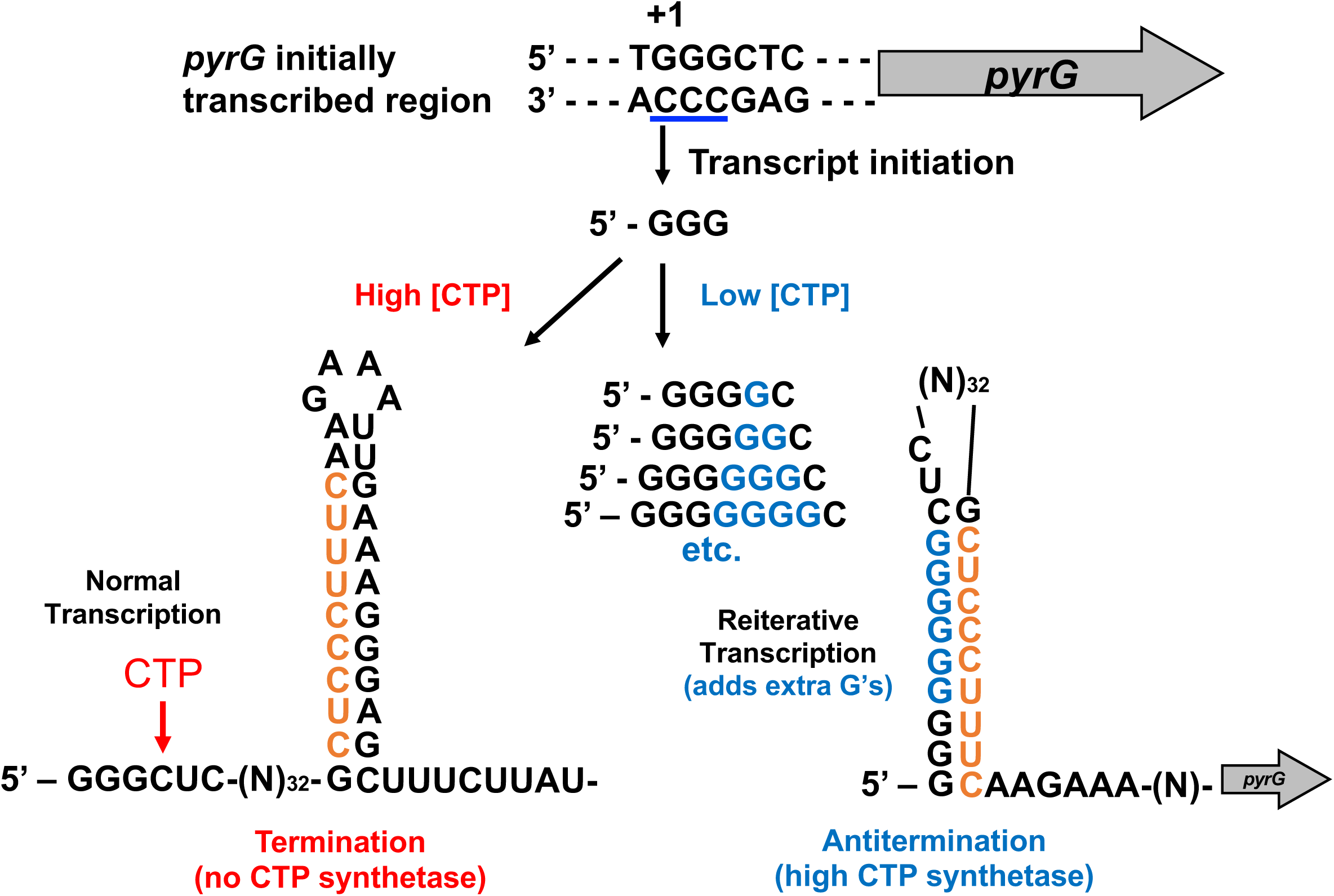
Model for CTP-mediated regulation of *pyrG* expression in *B. subtilis*. High CTP concentration allows canonical transcription by inserting CTP after the first three bases (5’-GGG-3’) are synthesized, resulting in the formation of the transcription termination hairpin, thereby eliminating *pyrG* expression (left). Low CTP concentration allows reiterative transcription which adds extra G bases to the nascent RNA right after 5’-GGG-3’ RNA is synthesized, resulting in the formation of the anti-termination hairpin, thereby allowing *pyrG* expression (right). Figure modified from reference (6).

**Fig. S2.**
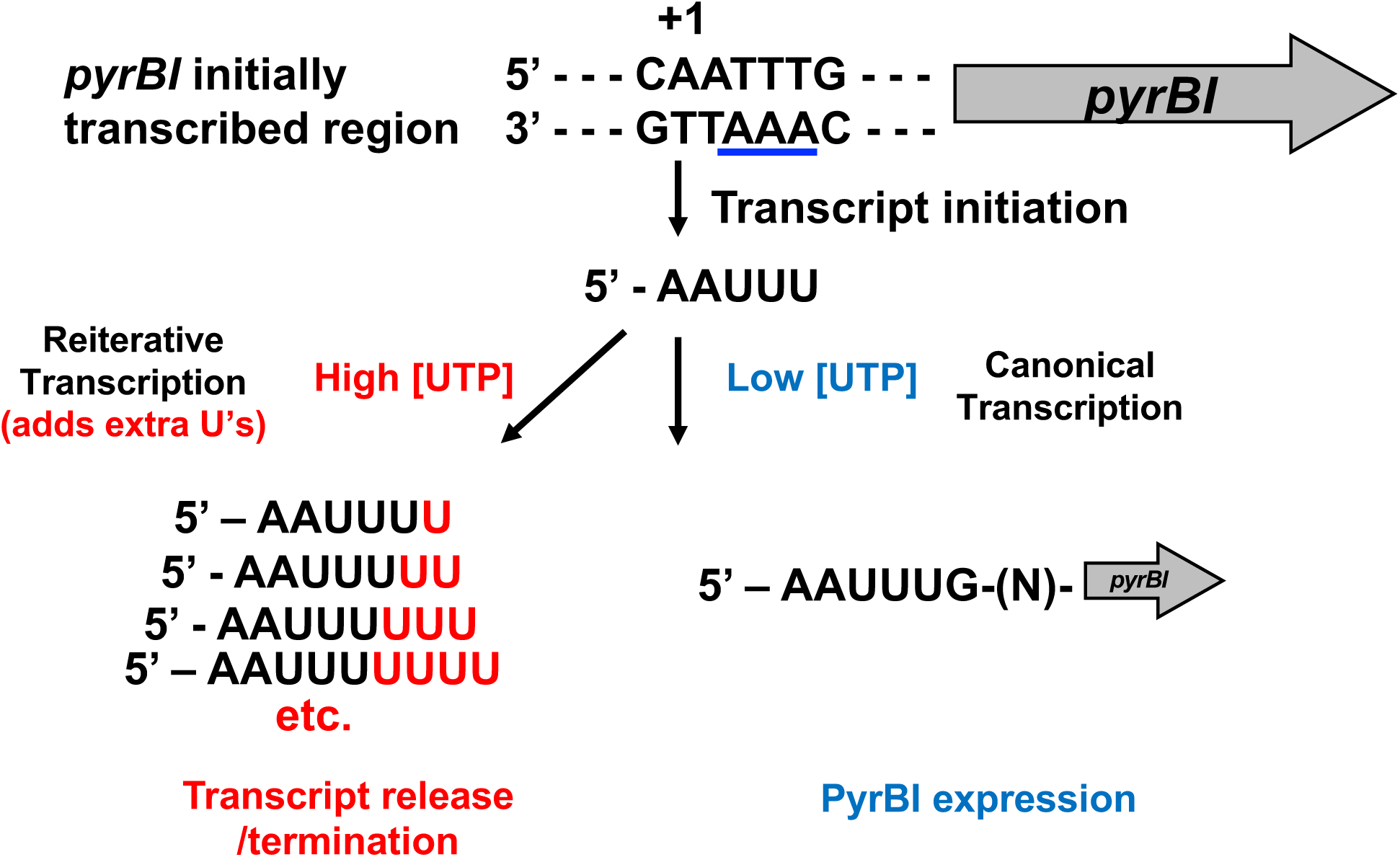
Model for UTP-mediated regulation of *pyrBI* expression in *E. coli*. High UTP concentration allows reiterative transcription that adds extra U bases to the nascent RNA (5’-AAUUU-3’). These transcripts are released from the transcript initiation complex, thereby *pyrBI* expression is reduced (left). Low UTP concentration allows canonical transcription by inserting GTP after the first five bases (5’-AAUUU-3’) are synthesized, thereby *pyrBI* expression is induced (right). Figure modified from reference (2).

**Fig. S3.**
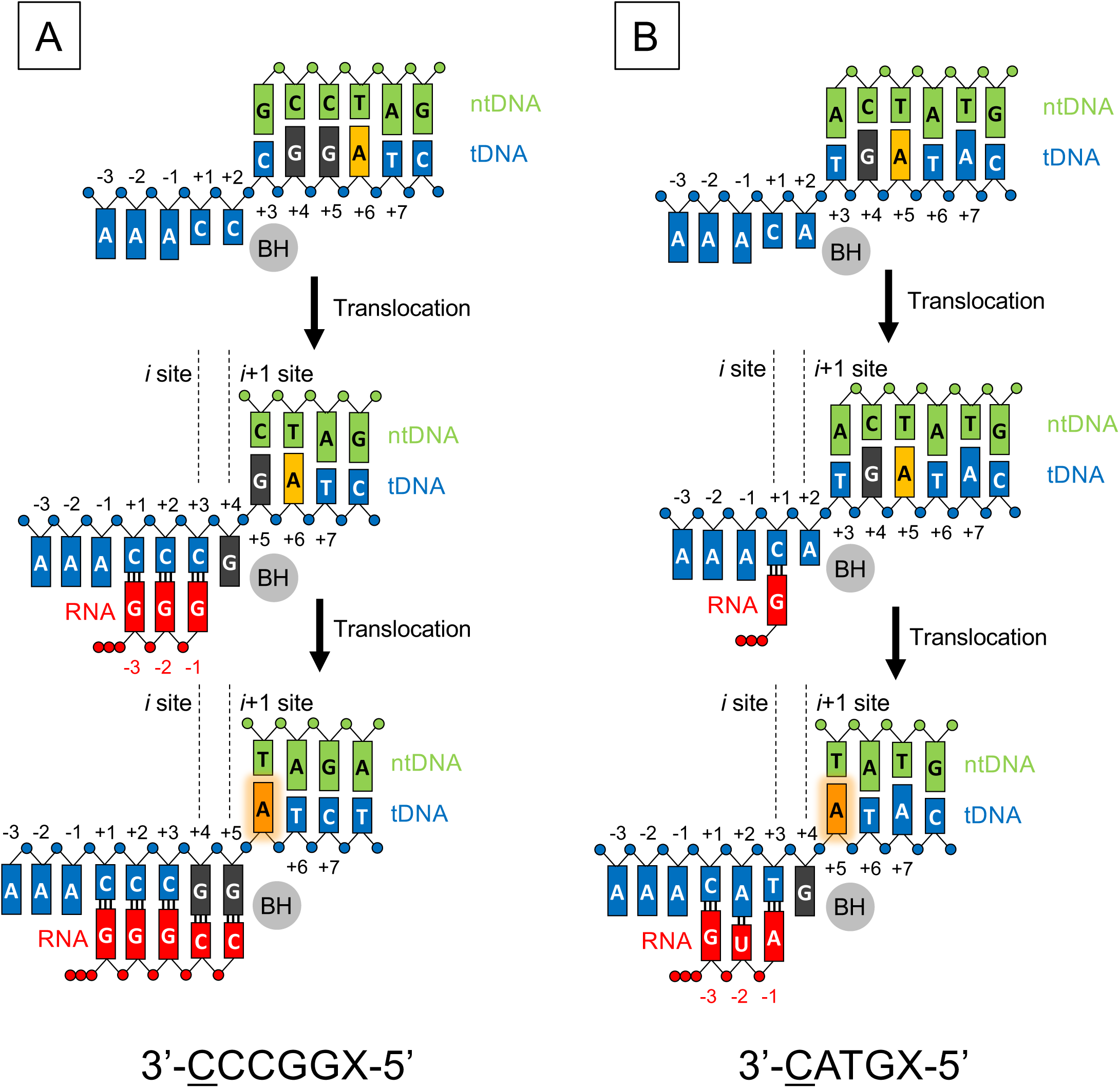
Reaction scheme of transcription at control DNAs. The DNA scaffold with tDNA sequence 3’-<u>C</u>CCGGX-5’ (transcription start site is underlined, X = 2-AP) contains *pyrG* initially transcribed region sequence but with an extra G base after +4G tDNA (left). The DNA scaffold with tDNA sequence 3’-<u>C</u>ATGX-5’ contains canonical transcription sequence (right).

**Fig. S4.**
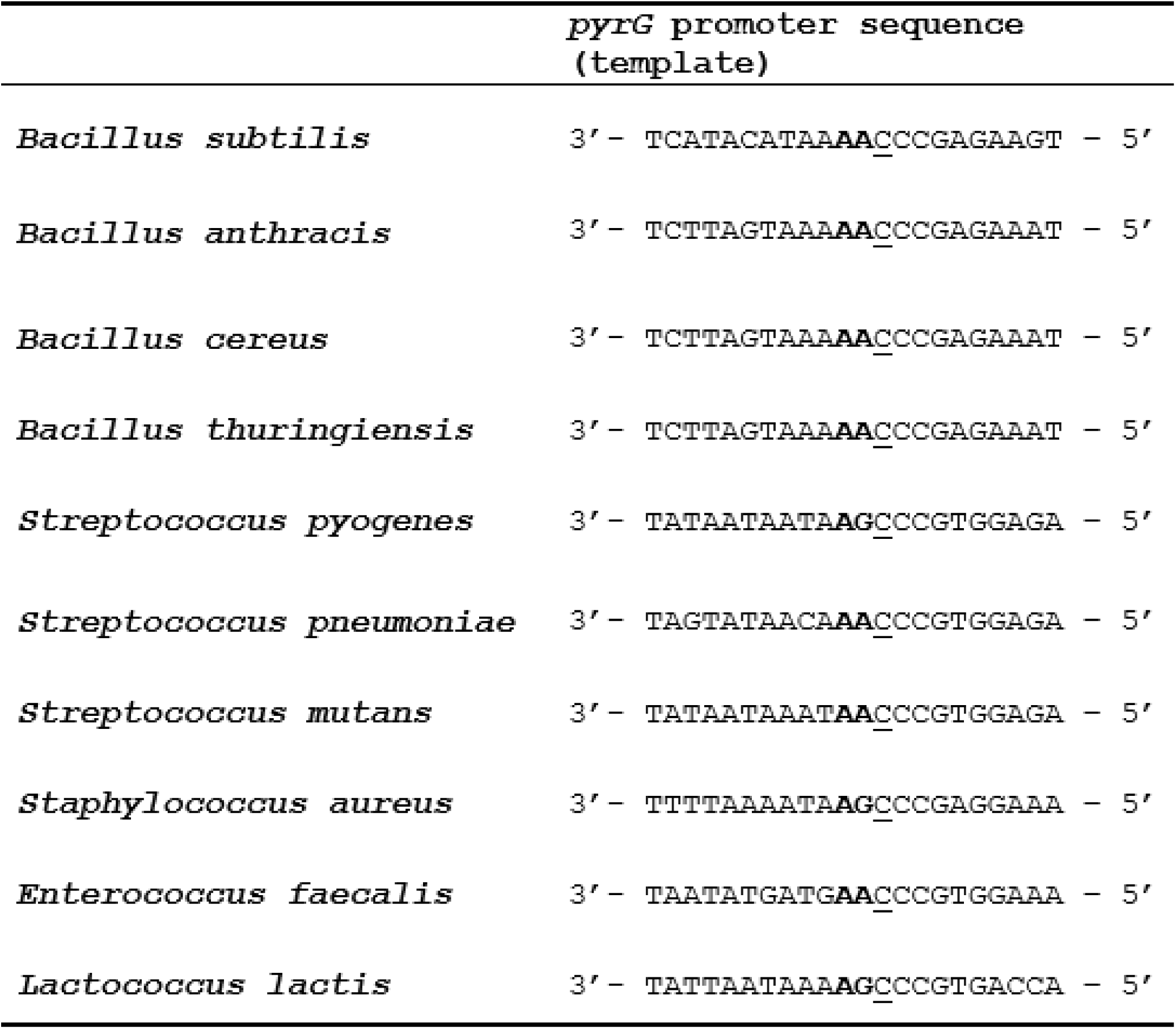
DNA sequences of the *pyrG* promoter. Template DNA sequences are shown. Transcription start sites are underlined and purine tracks from −2 to −1 positions are in bold.

**Fig. S5.**
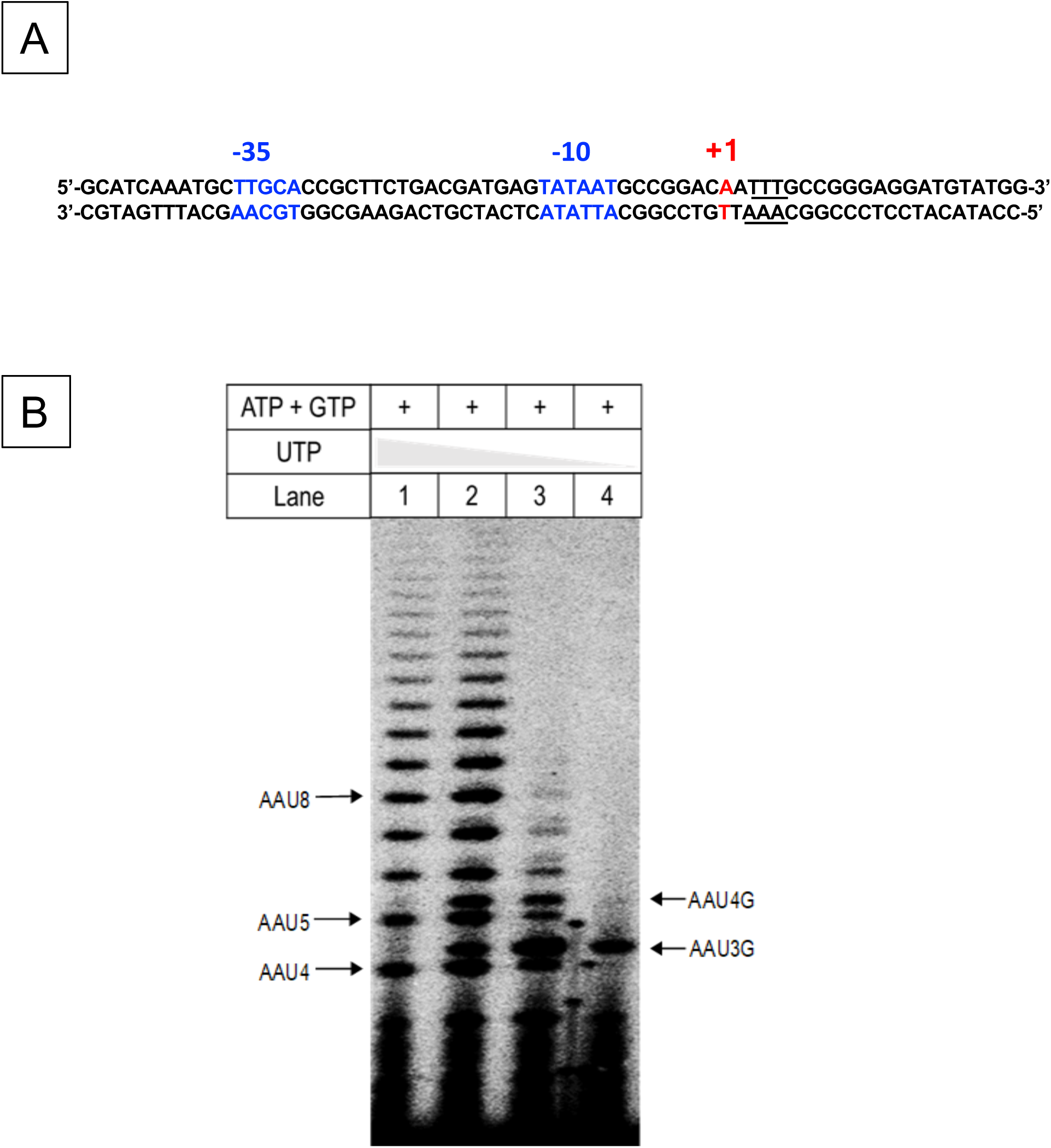
Characterization of reiterative transcription from the *pyrBI promoter*. A) DNA sequence of the *E. coli pyrBI* promoter region. The −35 and −10 regions (blue) and the +1 transcription start site (red) are indicated. Slippage prone sequence that allows reiterative transcription is underlined. B) *In vitro* transcription using DNA scaffold containing *pyrBI* ITR sequence. Transcription assay was performed in the presence of ATP (100 μM ATP and [γ-^32^p] ATP), 100 μM GTP, and different concentrations of UTP (5 mM, 1 mM, 100 μM, and 10μM).

